# Hepatitis B virus x (HBx) protein increases cellular OCT3/4 and MYC and facilitates cellular reprogramming

**DOI:** 10.1101/2023.08.11.552949

**Authors:** Madhusudana Girija Sanal, Sarita Gupta, Rahul Saha, Nisha Vats, Shiv Kumar Sarin

## Abstract

HBx is a multifunctional protein coded by the Hepatitis B virus which is involved in various cellular processes such as proliferation, cell survival/apoptosis, and histone methylation. HBx was reported to be associated with liver ‘cancer stem cells’. The stemness inducing properties of HBx could also facilitate the generation of pluripotent stem cells from somatic cells. It is well established that somatic cells can be reprogrammed to induced pluripotent stem cells (iPSC) using a cocktail of transcription factors called Yamanaka’s factors (OCT4, SOX2, KLF4, and MYC). The reprogramming process proceeds step-by-step with reprogramming factor chromatin interactions, transcription and chromatin states changing during transitions. HBx is a ‘broad spectrum trans-activator’ and therefore could facilitate these transitions. We electroporated low passage and high passage (difficult to reprogram) fibroblasts using Yamanaka’s factors (YF) with and without HBx and evaluated the reprogramming efficiency. We also investigated the tri-lineage and terminal differentiation potential of iPSC derived using HBx. We found, addition of HBx to YF, improves iPSC derivation, and increases the efficiency of iPSC generation from ‘difficult or hard-to-reprogram samples’ such as high passage/senescent fibroblasts. Furthermore, we show that HBx can substitute the key transcription factor MYC in the YF cocktail to generate iPSC. Our results have practical value in improving the efficiency of pluripotent stem cell derivation from ‘difficult to reprogram’ somatic cells in addition to providing some insights into the mechanisms of liver carcinogenesis in chronic hepatitis B. To conclude, HBx improves the reprogramming efficiency of YFs. HBx increases the cellular levels of OCT3/4 and MYC.

**Lay summary:** HBx is a protein made by hepatitis B virus. Yamanaka’s Factors (YF) are a mixture of ‘master-proteins’ which are used to convert adult cells to embryonic stem cells. We found that HBx protein can augment the efficiency of YF in generating these stem cells (concept summary figure).

## Introduction

Hepatitis B virus X protein (HBx) is 154 amino acids long (17kD) multifunctional protein produced encoded by hepatitis B virus (HBV). It is known to interfere with various cellular processes of the host such as transcription, signal transduction, cell cycle regulation, apoptosis and protein degradation(Kumar and Sarkar, n.d.; Minor and Slagle, 2014; Murakami, 2001). This multifunctional nature of HBx implies its role in altering several cellular mechanisms by hijacking the host cell homeostasis in favor of cancer initiation and evolution(Sivasudhan et al., 2022). HBx is known to activate the regulator transcription factor MYC (Terradillos et al., 1997) and prolongs its cellular levels by inhibiting its ubiquitination and proteasomal degradation(Kalra and Kumar, 2006). HBx is also involved in the regulation of gene expression through epigenetic mechanisms. For example, HBx relieves chromatin-mediated transcriptional repression of cccDNA of hepatitis B virus involving SETDB1 histone methyltransferase(Andrisani, 2021; Rivière et al., 2015). Wang et al, reported in 2017 that the HBx protein promotes the ‘stem-like’ properties of OV6+ cancer cells in hepatocellular carcinoma(Wang et al., 2017).

It is now well established that pluripotency can be induced in mature/well differentiated cells using Yamanaka’s factors – a cocktail of transcription factors OCT4, SOX2, KLF4 and c-MYC. Pluripotent stem cells generated using Yamanaka’s factors are known as ‘induced Pluripotent Stem Cells’ (iPSC)(Anonymous, 2016; Yamanaka, 2012). The use of MYC, a well-known proto-oncogene is often seen as a concern. In fact, each one of them (MYC, KLF4, SOX2 and OCT4), have been implicated in various cancers (Kim et al., 2015; Miller et al., 2012; van Schaijik et al., 2018; Wei et al., 2006) and all are essential for normal growth and development. The process of generating iPSCs with Yamanaka’s factors is not very efficient despite its relative simplicity(Brumbaugh and Hochedlinger, 2013; Winzi et al., 2014; Zhang et al., 2012). This is especially true for clinical samples obtained in small quantities where the number and quality of the cells could be poor (for example from a bio bank). The efficiency of the iPSC derivation decreases with increasing cell passage number and other parameters such as the age of the donor(Trokovic et al., 2015). Similarly, fibroblast samples from some individuals are very difficult to reprogram. Epigenetic modulators are known to increase the efficiency of iPSC derivation(Kang et al., 2014). Considering the role of HBx in cell proliferation, epigenetic pathways, protein stability, MYC activation and more importantly the fact that HBx imparts ‘cancer stem cell’ properties to transformed liver cells; we tested our hypothesis that addition of HBx in the Yamanaka’s reprogramming cocktail could enhance the efficiency of iPSC generation. We report here that HBx can not only augment the iPSC generation, but also could substitute MYC in the cocktail.

## Materials and Methods

### Cloning of Hbx

A synthetic sequence corresponding to 2A peptide from Thoseaasigna virus capsid protein and BspTI (AflII) restriction site was first cloned to pCXLE-EGFP (Plasmid #27082, Addgene, www.addgene.org) (Okita et al., 2011). The clones were verified by sequencing and subsequently digested with AflII (Thermo Fisher Scientific Cat. No. ER0831) to prepare the vector. Hbx insert was PCR amplified from pcDNA3-HBV(Gao et al., 2004) with Phusion DNA polymerase (New England Biolabs Cat. No. M0530S) according to the manufacturer’s protocol, gel purified using the GeneJet gel extraction kit (Thermo Fisher Scientific Cat. No. K0691) and digested with AflII. The vector and insert (1:2) were ligated using T4 ligase following the manufacturer’s instructions (Thermo Fisher Scientific Cat. No. EL0011). The cloned plasmid (here after denoted as pHBx) was verified by sequencing and by transfecting HEK293 cells and verifying the protein expression (Supplementary Figure/data-1a-c).

### Cell culture

HEK 293 cells were cultured in 10% fetal bovine serum (FBS), Dulbecco′s Modified Eagle′s Medium (DMEM) from CellClone, Genetix Biotech Asia, New Delhi, Cat.No.CCS-500-SA-U, supplemented with 1x penicillin (100U/ml) and streptomycin (100 µg/ml) (Hi-Media, Mumbai Cat. No. A018-5X100ML). At around 80% confluence, to validate the gene expression the plasmid (vide-supra) was transfected into HEK293 cells using PEI (Sigma-Aldrich/Merck Cat. No. 49553-93-7) following an optimized version of the standard protocol(Rajendra, 2018) with a DNA:PEI ratio of 1:3. The transfected cells were harvested by scrapping out on the third day for downstream applications.

A variety of fibroblast lines available in our lab that were grown following the established protocols from patient samples as well as healthy volunteers (approved by the institutional ethics committee whenever applicable or obtained from commercial sources such as biobanks) were used for the experiments (Rittié and Fisher, 2005). The derivation of fibroblasts from certain clinical samples resulted in poor quality fibroblasts, either a result of volume or quality of the specimen, because of age of the donors or other reasons (some of these fibroblasts showed the signs of senescence).

### Immunofluorescence/Immunohistochemistry/staining

Cells grown on the chamber slides were fixed with ice cold methanol (Sigma-Aldrich/Merck Cat. No. 34860) for 5 minutes at room temperature and then gently washed with phosphate buffered saline (PBS). Cells were incubated overnight in blocking buffer that contained the primary antibodies: OCT3/4, SSEA4, TRA-1-80, cytokeratin-18, HNF-4α, ASGPR1 (1:200; SC-5279, SC-21704, SC-21706, SC-32329, SC-374229, and SC-393849, Santa Cruz Biotech CA, USA), GATA4 (mAb #36966, Cell Signaling Technologies, MA, USA) antibody against human vimentin (pre-diluted, PathnSitu Biotechnologies CA, USA Cat # PM104), secondary goat anti-mouse Alexa Fluor® 568 (Abcam Cat. No. ab175473), anti-mouse Alexa Fluor-488 (Jax Immune Research, ME, USA Cat. No. 115-545-062). More details are provided in the CTAT supplementary table. After blocking with 5% FBS in TBST for 2 hours, cells were rinsed with TBST and incubated with appropriate secondary antibodies for 4 hours at room temperature in the dark. The nuclei were counterstained with DAPI. Images were acquired using fluorescence microscopes from Nikon Eclipse Ti-S or Evos M5000 (inbuilt grey scale camera and therefore all images are pseudocolored). Proprietary software associated with microscopes were used to acquire images. Inkscape 1.12 (inkscape.org), ImageJ (NIH, DC, USA), Paint (Microsoft, CA, USA), Irfanview 4.6 (www.irfanview.com) were used to view, analyze, adjust/ improve, label and layout the images and figures. Flow cytometry data was acquired by FACS Aria III and was analyzed by FACS data analyzed by Flowjo 10.4 (FlowJo LLC, Oregon, USA).

### Generation of iPSC

iPSCs were generated following established protocols(Anonymous, n.d.; Muthusamy et al., 2016; Ohnuki et al., 2009; Vaidyanath et al., 2020). Fibroblasts (high passage-over 15 passages and low passage -under 5 passages) were cultured on gelatin coated cell culture plates. On the fourth day, they were electroporated with Yamanaka’s plasmids (plasmid numbers 27078 (human *SOX2* and *KLF4*), 27080 (human *L-MYC*, *LIN28*), 27077 (human *OCT3/*4, shRNA against *TP53*) from Addgene, www.addgene.org) (Okita et al., 2011)plus or minus pHBx on Amaxa Nucleofector (Lonza, Cologne, France) using the manufacturer’s protocol for human dermal fibroblasts. We have also transfected the two groups of fibroblasts with pHBx alone. The electroporated cells were seeded on to ES qualified Matrigel™ (Corning, NY, USA product No. 354277) coated 6 well plates (∼10^5^ cells/cm^2^) with 10% FBS containing DMEM. The media was changed after 2 days to Essential 6 (Invitrogen Cat. No. A1516401) supplemented with b-FGF 20ng/mL (Peprotech US, Cat. No.100-18B) and the media was changed every two days till iPSC colonies started to appear.

### Picking of iPSC colonies and their maintenance

When colonies appeared (range: day 18 to day 30, from citing of the earliest ‘nidus’ appeared to the appearance of the last colony on any of the culture dishes), they were allowed to grow for 2 days to allow them to a size convenient for manual picking (when they reached about 200-300 µm size) and the picked colonies were allowed to grow larger and they ‘passaged’ by mechanical splitting into a new Matrigel coated plate containing mTeSR™1 media with its supplement(Anonymous, n.d.) and maintained as suggested by the manufacturer (StemCell Technologies, Canada Cat. No. #85850). Subsequent passages were done by enzymatic dissociation (Accutase®, ThermoFisher Scientific, MA, USA, Cat. No. A11105-01) when the colonies reached 50-60% confluence.

### Western Blot

Whole-cell protein extracts were prepared from scraped out HEK 293, fibroblasts or iPSC culture plates and extracted using RIPA lysis and extraction buffer. Extracts were run on 10% SDS-PAGE and transferred to a polyvinylidene difluoride membrane using a transfer apparatus following the standard protocols (Bio-Rad). After incubation with 5% nonfat milk in TBST (10 mM Tris, pH 8.0, 150 mM NaCl, 0.5% Tween 20) for 60 min, the membrane was washed once with TBST and incubated with rabbit antibodies against human OCT-3/4 antibody (Santa Cruz Biotechnology Inc., CA, Cat. No. SC-5279), 1: 1000 dilution; mouse anti-HBx antibody (Thermo Fisher Scientific, MA, Cat. No. MA1-081), 1:1000, overnight incubation at 4°C. All blots were standardized with GAPDH as the internal reference protein. Mouse anti-GAPDH antibody (Proteintech Cat. No.60004-1-Ig) 1:10000, dilution 1:1000 at 4°C overnight. The membrane was washed with TBST buffer and incubated with a 1:5000 dilution of horseradish peroxidase-conjugated anti-rabbit (Santa Cruz, CA, Cat# SC-2004)/anti-mouse antibodies (Santa Cruz, CA, Cat. #SC-2005) for 2 h at room temperature. Blots were washed with TBST four times and developed with the ECL system (Bio-Rad, US Cat. #170-5060), according to the manufacturer’s protocols. Chemiluminescence imaging was done using iBright 1500 (Thermo Fisher Scientific, MA, USA).

### Real-time polymerase chain reaction

Total RNA was isolated using NucleoZOL (Takara, Shiga, Japan, Cat. No. 740404.200) following the manufacturer’s instruction. cDNA was prepared from (deoxyribonuclease treated) total RNA using RevertAid Reverse Transcriptase (Thermo Fisher Scientific, MA, Cat. No. EP0441) following the manufacturer’s instructions. Real Time quantitative PCR (RT qPCR) was done using specific oligonucleotide primers targeting the genes of interest (supplementary data, table 1), denaturing temperature 95°C, annealing temperature 60°C, extension temperature 72C in triplicates and two repeats, using GoTaq® qPCR Master Mix (Promega Cat. No. A6001) following ‘manufacturer’s instructions on a Veriti Thermo Cycler (Applied Biosystems, MA, USA). The data was acquired using the software associated with the same machine (ViiA7 V1.2) and relative quantification with respect to the housekeeping genes was calculated using the by 2 (–ΔΔCt) method.

### Screening of iPSC colonies

To identify and estimate the efficiency of iPSC colony formation from fibroblasts, the six well plates were stained for the activity of alkaline phosphatase as iPSC colonies are known to produce alkaline phosphatase(Vaidyanath et al., 2020). We used BCIP/NBT tablets (SigmaFast Sigma-Aldrich/Merck Cat. No. B5655) as a substrate for alkaline phosphatase which produces a deep purple-black precipitate when alkaline phosphatase is present. The plates were imaged after three weeks of transfection. To estimate the efficiency of iPSC colony formation from fibroblasts, purple-black spots larger than 50 microns were counted in four fields (4 x objective).

### Characterization of iPSC colonies

The picked colonies were passaged over 15 times to check whether the colony morphologies retain the acceptable morphology under light microscopic examination. The iPSC colonies were tested for the expression of alkaline phosphatase (Invitrogen, USA Cat. No. A14353), OCT3/4, SSEA4 and TRA 1-81. Differentiation of iPSC: iPSC cells were aggregated to form embryoid bodies which were differentiated to ectodermal, endodermal and mesodermal lineages following the established protocols. The germ layers were evaluated using specific markers by immunofluorescence microscopy (Ohnuki et al., 2009),(Muthusamy et al., 2014),(Baghbaderani et al., 2016). iPSCs were subsequently differentiated to ‘hepatocyte like cells’ following a modified version of the protocol described by Basma et al.(Basma et al., 2009).

Genomic DNA was isolated from iPSC using DNEasy blood and tissue kit (Qiagen, Germany Cat. No. 69504). DNA Polymerase Chain Reaction (PCR): to verify persistence/integration of HBx DNA in iPSC lines: A PCR was done investigate the persistence (as an episome) or integration (genomic) of the transfected HBx DNA in the iPSC derived from the fibroblasts transfected. The same forward and reverse primes (Ta=60C) were used for the PCR reaction (Supplementary table 1). Taq DNA polymerase was used for the PCR reaction following the manufacturer’s protocol. The PCR products were resolved on a 1% agarose gel in TAE buffer(Voytas, 2001).

### Karyotyping

Briefly when the plates (fibroblasts/iPSC) reached 80% confluence, cells were treated with colcemid for 30 minutes at a final concentration of 0.1 µg/ml with media. The cells were harvested by trypsinization, washed in PBS, pelleted down by centrifugation at 200g. The cells were resuspended in 0.075M potassium chloride and incubated at 37 °C for 20 min in a volume of 2 ml. Cells were pelleted down. Eight drops of fresh ice-cold fixative (3:1, methanol: acetic acid) was added to suspend the cells. The volume was brought up to 0.5mL with the fixative solution and was incubated for 15 minutes at room temperature. This was repeated twice. Following a last spin, the cells were resuspended in 200–300 μl of fresh fixative, and dropped onto clean slides, air dried and a proportion of metaphase spreads were imaged and assembled(Anonymous, n.d.).

### Cell cycle, proliferation, and apoptosis analysis

We performed cell cycle analysis and apoptosis assay using flow cytometry with propidium iodide (SRL, Mumbai, India, Cat No. 11195) a standard nuclear stain(Crowley et al., 2016; Riccardi and Nicoletti, 2006). The cell proliferation assay was done using the standard MTT assay (Sigma-Aldrich/Merck, Catalog No. 11465007001). MTT assay was performed on each well following the manufacturer’s protocol.

Briefly, equal numbers of cells were seeded on six well plates. The test wells were transfected with pHBx and control was transfected with GFP expressing plasmid (Plasmid #27082, Addgene). After three days post transfection transfected cells were treated with low dose puromycin (0.5 μg/ml) for 6 hours. Untransfected cells with and without puromycin treatment, transfected cells with and without puromycin treatment were subjected to flowcytometry with propidium iodide to estimate apoptosis in each experimental group. pHBx transfected and control cells were also used for cell cycle analysis using propidium iodine.

### Statistics

Data are reported as meansL±LSD whenever applicable from at least three independent experiments. Student’s t-test was performed to compare the mean between two groups and one-way ANOVA when more than two groups were compared. Numerical data was saved and processed using Microsoft Excel (2013) Washington, United States and Graphpad Prism 7.04, Graphstats Technologies USA.

## Results

### HBx improves the generation of iPSC

To verify whether the HBx expression vector (pHBx) is competent to express the viral protein HBx in mammalian cells, we transiently transfected HEK293 cells and fibroblasts, and total cell lysate was probed by western blot using an anti HBx specific antibody as detailed. Fig. 1A demonstrates that the 17kD specific band in cell extracts from pHBx transfected cells is absent in mock transfected cells.

**Figure-1.**
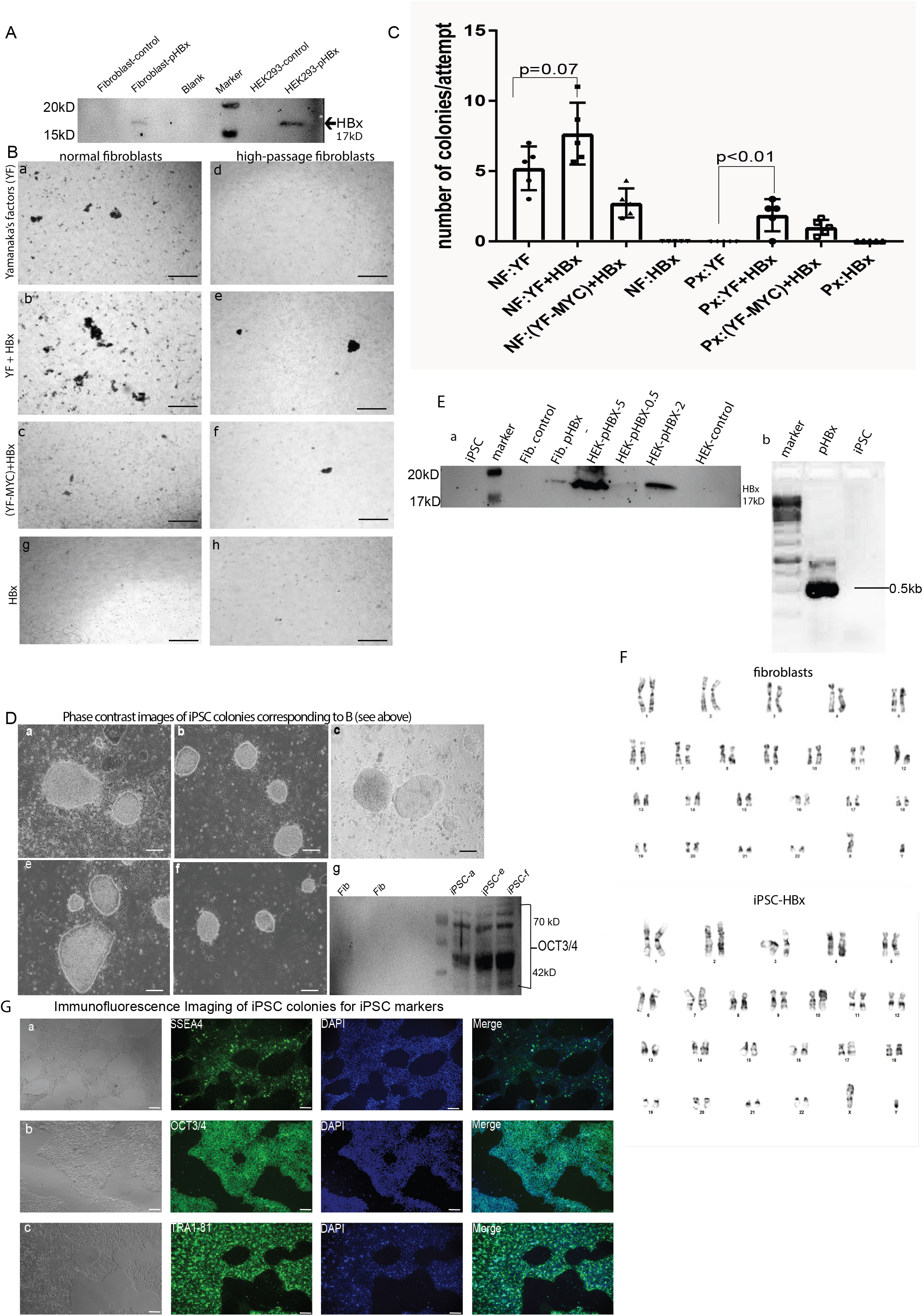
A. Western blot (qualitative) showing the expression of HBx (17kD) protein from total protein isolate from human fibroblasts and HEK293 cells (which were transfected with HBx plasmid (pHBx) and untransfected control). Full length blot is presented in supplementary figure-1-A. B. Human dermal fibroblasts (standard low passage normal fibroblasts marked NF-a, b, c / high passage fibroblasts marked Px-d, e, f) were transfected with Yamanaka’s factors (YF: a and d); Yamanaka’s factors along with HBx expression vector-pHBx (YF+pHBx : b and e); and Yamanaka’s factors where *MYC* was substituted with HBx (YF-MYC+pHBx : c and f). Normal fibroblasts (g) and high passage fibroblasts (h) were transfected with pHBx alone. The cell culture plates were screened for alkaline phosphatase activity using BCIP-a substrate of alkaline phosphatase which forms purple-black precipitate when acted upon by the enzyme (black spots on grey-scale images) Scale bar: 400µm. C. The number of colonies were counted under low power objective (4x) in four fields and the average number of colonies were plotted in each group on different attempts (‘n’ represents the number of replicates in each category). A t-test was used to calculate the statistical significance. D. iPSC colonies derived with pHBx showed similar morphology as they were picked, mechanically split and grown on Matrigel™ coated plates (a, b, c, e, f corresponds to a, b, c, e, f in figure-1B) compared to iPSC derived using the standard YF. Scale bar: 100µm. g. Western blot (qualitative) was performed using total protein extracted from iPSC colonies derived using HBx showed that these colonies were positive for OCT3/4 a key transcription factor responsible for pluripotency. Full length western blots are provided in supplementary figure-1-D-g. **E.** Western blot demonstrating the loss of HBx expression in iPSC colonies derived with the help of HBx. Equal amounts of proteins were loaded. (a). Full length blot is presented in supplementary figure-1-E-a. Genomic DNA from these iPSC colonies were tested for the presence of HBx sequences derived from pHBx. No bands corresponding to HBx could be detected by PCR using HBx sequence specific primers (b). Uncut gel image is provided in supplementary figure-1-E-b. F. Karyotyping was to done to see any gross chromosomal anomalies in iPSC derived using HBx. The karyotype of pHBx derived iPSC was similar to the karyotype of human fibroblasts which were used to derive them. G. Immuno-fluorescence imaging demonstrating the conventional markers of pluripotent stem cells. 80% confluent pHBx derived iPSC colonies grown on Matrigel™ coated surface were imaged. These colonies were positive for OCT3/4 (nuclear marker), SSEA4 and TRA-1-60 (surface markers). Adjustments for individual channels were necessary on merged images. Scale bar: 100µm

**Figure-2.**
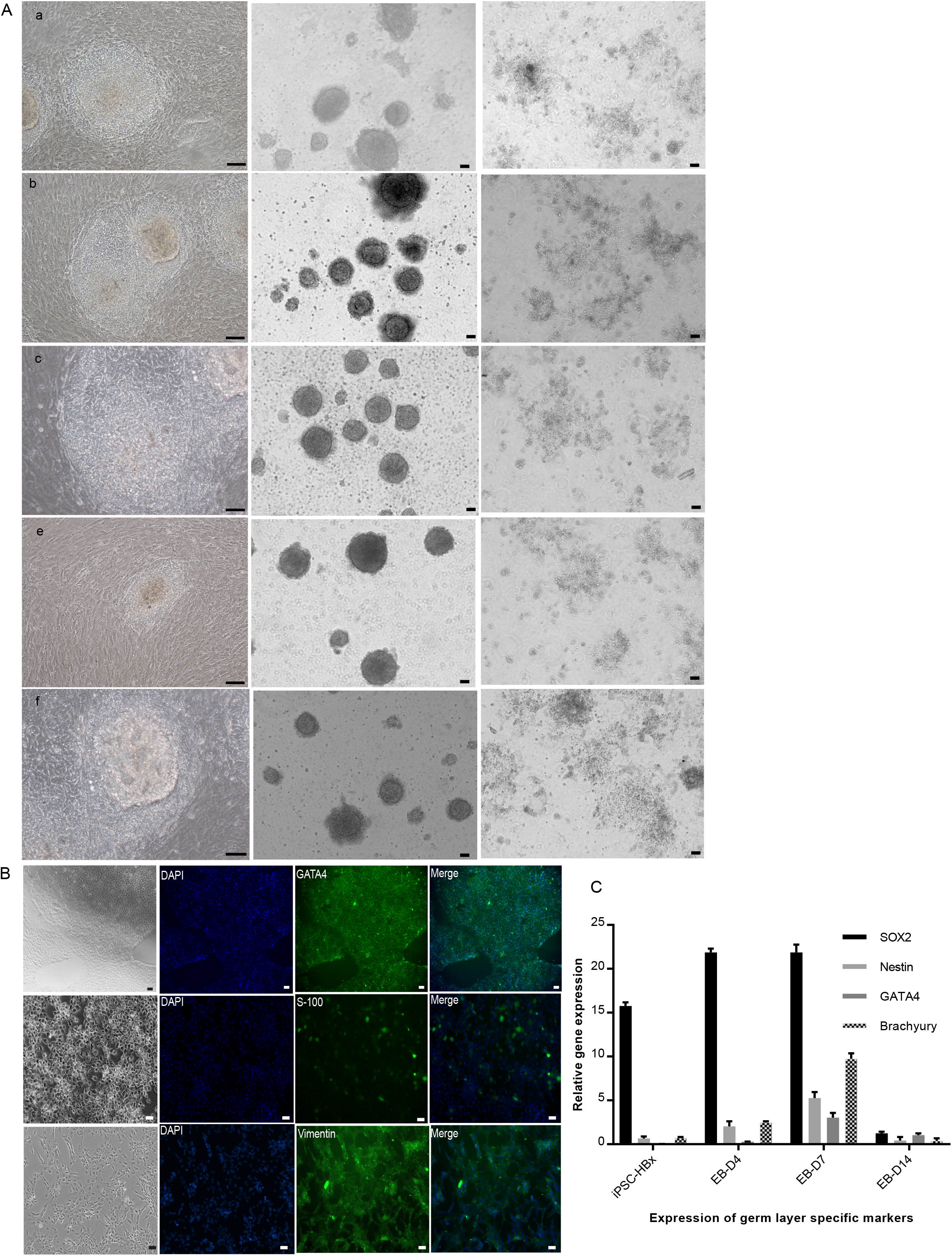
A. The iPSC derived using the standard YF or YF in combination with pHBx were stable and showed similar morphologies after fifteen passages. iPSC derived using pHBx were indistinguishable from standard iPSC. They showed central differentiation, they formed embryoid bodies (EBs) and started differentiation in the absence of FGF-2 in the maintenance media (labeled a, b, c, e, f as in figure-1D). B. Immuno-fluorescence imaging demonstrating the capacity of iPSC derived using HBx to form all the three germ layers: the pHBx derived iPSC upon differentiation to EBs expressed the markers of all the three germ layers –endoderm (GATA4), ectoderm (S100) and mesoderm (vimentin). Scale bar: 50µm C. The qPCR showed the changes in the expression of SOX2 (a key transcription factor present in iPSC, neural and endoderm progenitors, nestin (ectodermal marker), GATA4 (endodermal marker) and brachyury (mesodermal marker) were consistent with differentiating EBs. Adjustments for individual channels were necessary on merged images.

Human dermal fibroblasts passage 3 (HDF-P3) were transfected with Yamanaka’s factors containing plasmids in the absence (YF) or presence of pHBx (YF+HBX) as described in Okita K et al 2011(Okita et al., 2011). Colonies started appearing by day 20 and we screened the plates for the expression of alkaline phosphatase, a commonly marker used for screening of iPSC colonies, using NBT, an alkaline phosphatase substrate which gives a dark colored precipitate. Fig. 1B shows that addition of HBx resulted in an increased number of alkaline phosphatase positive cell clusters (compare 1A and B) suggesting an improvement in iPSC derivation from fibroblasts.

### HBx could replace c-Myc in Yamanaka’s cocktail

MYC has been reported to be an oncogene thereby developing a system lacking MYC would be advantageous. Considering the capacity of HBx to induce stemness in liver cancer stem cells, its ability to potentiate and increase the stability of endogenous MYC, we then investigated whether HBx could substitute for MYC in the Yamanaka cocktail. HDF-P3 cells were transfected with Yamanaka’s factors without MYC but supplemented with HBx expression vector. Fig. 1B, panel ‘a’ and ‘c’, shows that fibroblasts transfected with the Yamanaka cocktail or the Yamanaka cocktail in which c-MYC was substituted with HBx produced iPSC colonies with similar efficiency (Figure-1C). Thus, our data shows that HBx can substitute for c-MYC in Yamanaka’s factors.

It is well known that samples from high passage fibroblasts (fibroblasts maintained in culture for a long time and undergone multiple rounds of cell divisions) are difficult to reprogram(Trokovic et al., 2015). To test whether introduction of HBx in Yamanaka’s cocktail can improve reprogramming, we transfected high passage fibroblasts (HDF-Px) which failed to give iPSC colonies under standard conditions (Figure-1B-d) in our previous experiments with Yamanaka’s factors in the presence or absence of pHBx. We observed that these ‘difficult to reprogram’ cells produced iPSC colonies although the number of colonies were comparatively less (Figure-1B-e, f). The results are summarized in Figure-1C. In short, although addition of HBx gave more iPSC colonies, the results were not statistically significant (p=0.56). However, the difference was significant in high passage fibroblasts (p=0.01). We picked these colonies for further propagation and these colonies appeared to have the characteristic human iPSC/ES morphology under light microscope (Figure-1D-a-c and e which corresponds to a-b and e in figure-1B) and iPSC colonies were positive for OCT3/4, a key marker of pluripotent stem cells (western blot figure-1D-g).

### HBx sequence do not integrate to iPSC lines

Integration of HBx sequence into iPSC genome or persistence of HBx plasmid is a concern because HBx is associated with hepatocellular carcinoma. Genomic integration is a theoretical possibility associated with any transfection which involves DNA. We performed a Western Blot to see whether the iPSC generated out of pHBx transfection continues to express HBx protein in the event of plasmid integration or persistence. Total cell lysate from iPSCs at fifth passage was analyzed by western blot. Our results suggest that the plasmid gets lost as HBx expression was no longer detectable in cells which are reprogrammed to iPSC (Figure-1Ea).

Next, we checked whether any integrated HBx sequence can be detected by PCR in the iPSC. Using HBx sequence specific primers and genomic DNA isolated from iPSC lines as the template, after 34 cycles of PCR we could not detect any amplicon of the expected size on agarose gel electrophoresis compared to the positive control (pHBx plasmid) (Figure-1Eb). The iPSC derived using p-HBx and the fibroblasts used to derive the iPSC were karyotyped and compared to exclude any abnormalities incorporated during the reprogramming process. Figure-1F shows that the two sets are indistinguishable. We have also performed an RNASeq analysis of the iPSC derived using HBx and the preliminary analysis showed no significant difference with reference to the control (data not shown).

### Characterization of iPSC lines

We next characterized the colonies for the expression of the markers of pluripotency-OCT3/4, SSEA4 and TRA 1-81 by immunofluorescence microscopy (Figure-1G a-c). iPSCs were maintained the typical morphology under light microscope after 15 passages; they showed characteristic center differentiation (Figure-2A, right column), formed embryoid bodies (middle column) and started differentiation when FGF-2 was withdrawn from the culture medium (left column). iPSCs also differentiated into all the three germ layers and accordingly expressed markers specific for each distinct germ layer (Figure-2B). We analyzed the kinetics of expression of SOX-2, nestin, GATA-4 and brachyury by RT qPCR (Figure-2C). The expression of GATA4, nestin and brachury increased from day 0 to day 7 and decreased again by day 14. We further differentiated the iPSCs to hepatocyte like cells (i-Heps). These cells were positive for markers characteristic of hepatocytes-nuclear factors (FOXA2 and HNF4A), cytoplasmic factors (albumin, α-fetoprotein and coagulation factor-9), cytoskeletal protein (CK18) and surface protein ASGPR1 (Figure 3 a-g).

**Figure-3.**
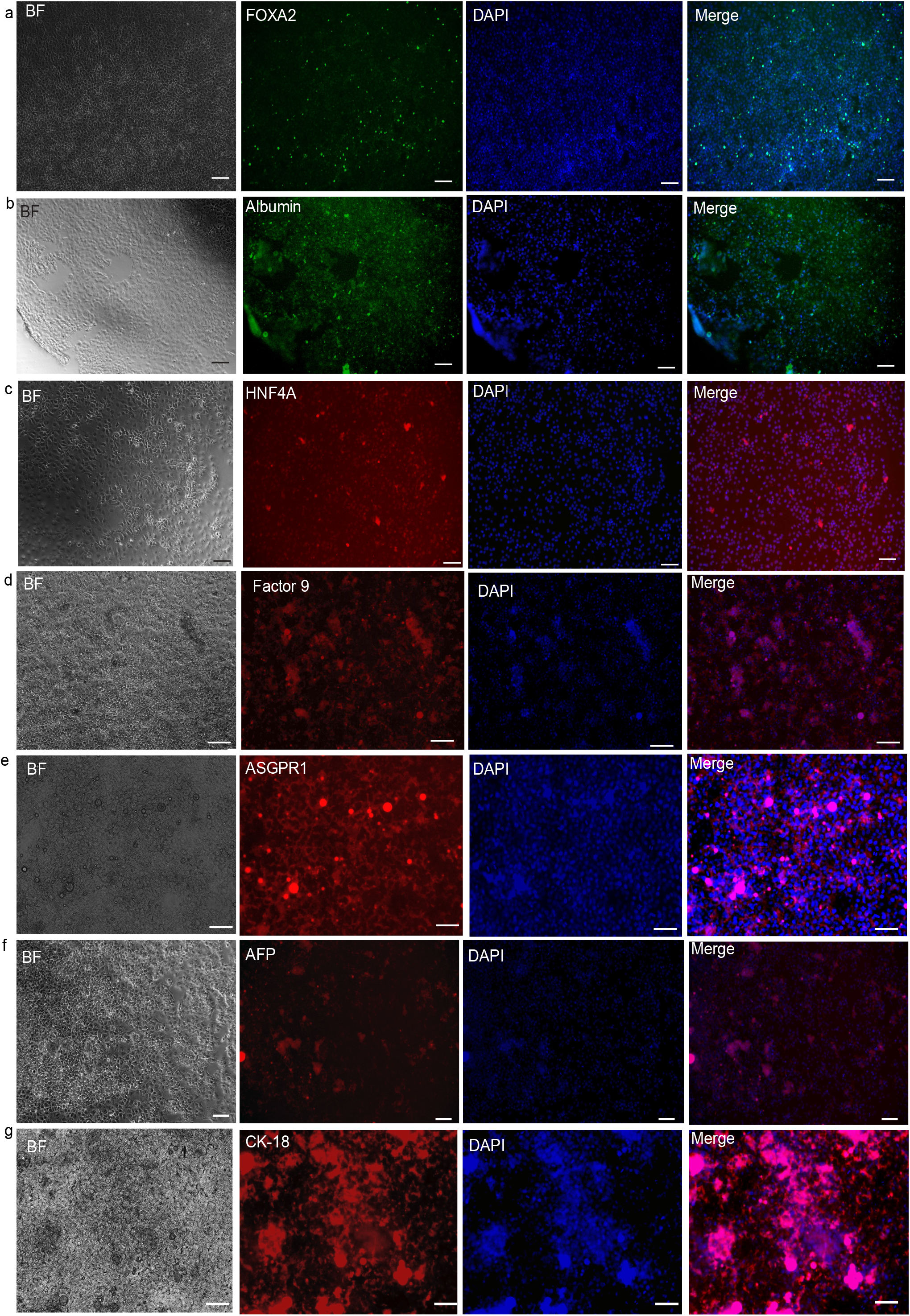
Bright field (BF) and immuno-fluorescence imaging of hepatocyte like cells showing lineage specific markers. The pHBx derived iPSC were differentiated to hepatocyte lineage to demonstrate their terminal differentiation potential. These terminally differentiated cells were positive for hepatocyte specific markers such as FOAX2 (a), HNF4A (c) (nuclear receptors), albumin (b), blood clotting factor 9 (d), AFP (f) (secreted proteins), ASGPR1 (e) (membrane protein), and CK18 (g) (cytoskeletal protein). Adjustments for individual channels were necessary on merged images. The iPSC control images are provided in the supplement data 1d. Scale bar: 100 µm

### Markers of stemness are lost upon differentiation

An important determinant of iPSC quality is the ability to undergo complete differentiation. Therefore, one potential concern could be the persistent expression of stem cell markers after induction of differentiation or incomplete differentiation. Incomplete differentiation raises the possibility of genetic instability and cancer if injected into animals (precluding clinical applications). Our immunofluorescence imaging showed loss of Oct4 with an increase in GATA4 expression (Figure-4A). RT qPCR analysis of mRNA from standard iPSCs, iPSCs generated using HBx, embryoid bodies (EBs) and iHeps (generated from HBx iPSC) showed that no differentiated cells, EBs or iHeps, expressed pluripotent stem cell markers OCT3/4 or NANOG (Figure-4B).

**Figure-4.**
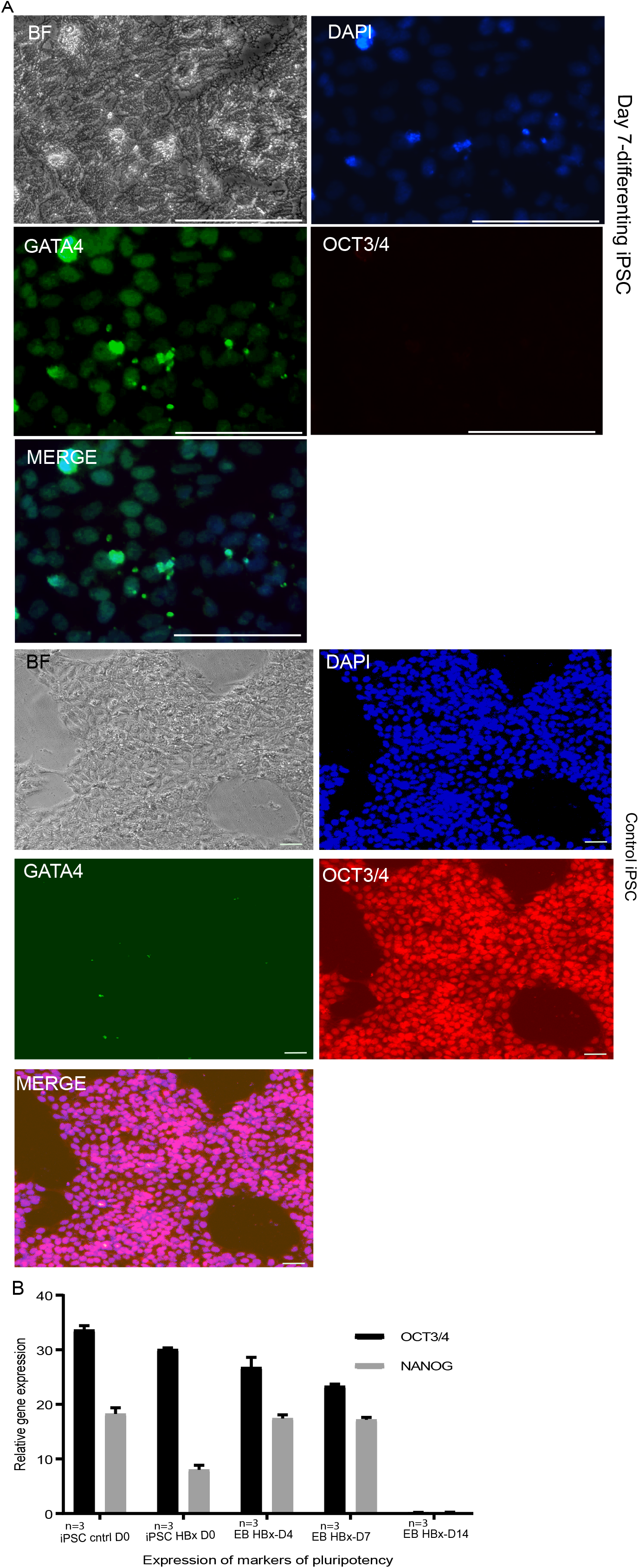
A. Immuno-fluorescence imaging demonstrating the loss of OCT3/4 as iPSCs underwent differentiation (above). The pHBx derived iPSC upon differentiation lost the expression of OCT3/4 a marker of pluripotency while the differentiated cells acquired GATA4 positivity. Below: Control images of iPSC cells. Scale bar: 50µm B. The qPCR analysis showed progressive down regulation of stemness markers OCT3/4 and NANOG confirming the differentiation. The difference between groups were statistically significant when iPSC, day 4 and day 7 were compared with day 14 for both OCT3/4 and NANOG (p<0.05, calculated by ANOVA).

### HBx increased the expression of MYC, OCT3/4 and Ki67

We investigated whether the property of pHBx to substitute for MYC in the Yamanaks’s cocktail could depend on its ability to induce MYC or to prolong its half-life. To check whether HBx increases the cellular levels of MYC we transfected HEK293 cells with pHBx and we found that there is an increase in the levels of MYC on western blot. Considering the promiscuous nature of HBx as a transactivator, we decided to check whether HBx could also increase OCT3/4. Our western blots showed an increased level of OCT3/4 in the transfected cells. We further checked whether here is a concomitant increase in the levels of Ki67, a nuclear protein that is associated with cellular proliferation and as expected we found an increase in the Ki67 levels in pHBx transfected cells compared to the control (Figure 5A). These results are summarized in the densitometric plots of western blots (Figure-5B). We used HEK cell line for these experiments because they are much easier to transfect compared to human dermal fibroblasts (but we confirmed the expression of HBx in both fibroblasts and HEK in the beginning of the reprogramming experiment. Refer figure-1A). We assumed that the mechanism could be the same for fibroblasts (which may not be the case). This is a major drawback of this study. Cell death at various stages adversely affects the efficiency of iPSC generation. It is well known that during the reprogramming process lots of cells undergo apoptosis and decrease in cell death would have a positive effect in reprogramming efficiency. Cell death ranges from apoptosis of fibroblasts at the time of transfection to apoptotic events during the reprogramming process. To check this, we did a flow cytometric analysis of pHBx transfected cell which revealed that cells transfected with pHBx, cell death decreased significantly (about 12% in control versus ∼6% in transfected cells, p<0.05). This effect was especially pronounced when cells were exposed to stress (low concentration puromycin treatment for 24h). About 47% cells died in puromycin treated plates compared to ∼35% in pHBx transfected cells treated with puromycin (p<0.05) (Figure-5C). More cells moved to proliferation-the cell in G2/G1 was more in pHBx transfected cells (G2:13%) compared to control (G2:8.7%; Figure-5D). An MTT assay was performed (third day post-transfection) to assess the metabolic activity but the difference between pHBx transfected and control cells were not statistically significant (Figure-5E).

**Figure-5.**
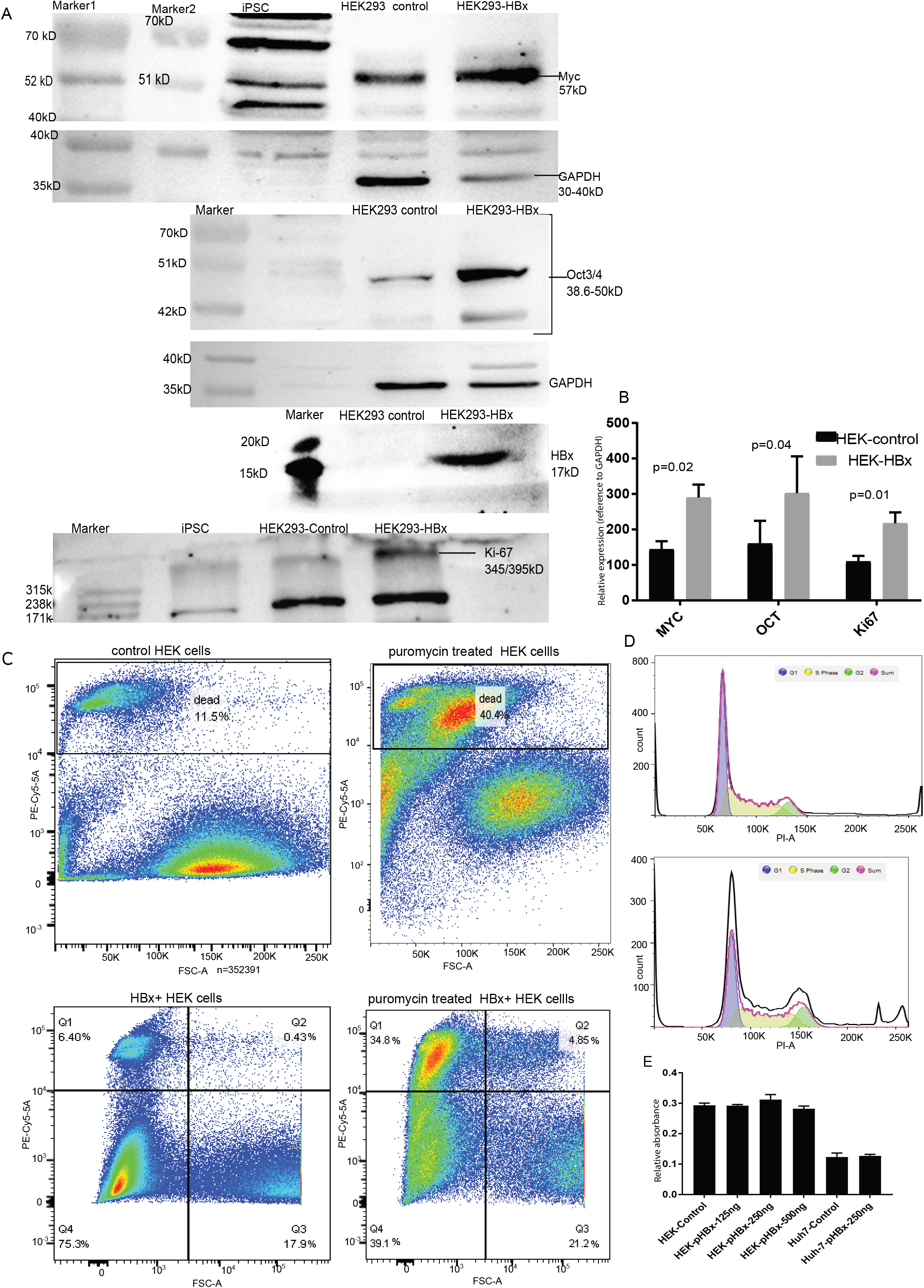
A. Western blot demonstrates the potential mechanisms were by HBx protein improves the reprogramming process: Transfection of pHBx resulted in resulted in increased MYC, OCT3/4 and Ki67. Equal amounts of total protein of pHBx transfected and untransfected (control) HEK293 cells were loaded (except for the iPSC total protein which was run to serve as control only to ascertain the position of OCT3/4 band). GAPDH blots are provided for comparison. The Full length blots are presented in supplementary figure-5-A (labeled row-1-6). B. The densitometric plot of the relative band intensities with reference to GAPDH. The difference in MYC, OCT and Ki67 expression in pHBx transfected cells were statistically significant compared to the untransfected control. A paired t-test was used to compute the statistical significance. **C.** Flowcytometric analysis based on propidium iodide (PE-Cy5-5, 690/50 band pass filter) to verify cell death and cell division: pBx transfection resulted in improved cell survival. This effect was more pronounced under a stressor, puromycin. Fewer pHBx transfected cells died upon treatment with low dose puromycin. D. Flowcytometry for DNA content using propidium iodide suggests more cells in G2 phase in pBx transfected cells. E. MTT assay: There was no significant difference between the untransfected control cells versus pHBx transfected cells suggesting similar levels of metabolic activity (NADPH) in transfected and untransfected wells. The untransfected control was compared with each test sample independently (paired t-test).

## Discussion

HBx is a multifunctional protein coded by the hepatitis B virus which is critical for the survival of the virus within human body. It is possible that HBx helps the virus to ‘reprogram’ the epigenetic landscape of hepatocytes to facilitate its replication, and in this process, cells mutate to form hepatocellular carcinoma. Upon induction of the HBx gene, resting cells begin to enter the S-phase of the cell cycle(Koike, 1995).Promotion of cell division is important for viral survival and infectivity. HBx is also known to inhibit the ubiquitination and proteasomal degradation of c-Myc(Kumar and Sarkar, n.d.).The HBx also induces growth factors, activates MAP and Src kinases, inhibits apoptosis by multiple mechanisms including inhibition of p53(Kumar and Sarkar, n.d.). HBx is known to be a ‘broad spectrum trans activator’ with the ability to regulate all three classes of promoters(Kumar and Sarkar, n.d.). Furthermore, HBx was reported to induce ‘cancer stem cells’ in hepatocellular carcinoma^10^. Considering these facts, we presumed that the addition of pHBx could increase iPSC generation in cells which are difficult to reprogram such as cells isolated from old patients, primary cells which are poorly characterized, ‘cell lines’ derived from very small quantity or poor-quality tissue samples (for example from rare diseases). Therefore, our findings could be useful in iPSC generation facilities such as biobanks or iPSC repositories.

Our studies do not resolve the mechanism by which HBx improves the outcome of iPSC generation. One possibility is that HBx increases the amount of endogenous MYC in the transfected fibroblasts by prolonging the half of intracellular MYC. Our experiments demonstrated increased levels of MYC upon HBx expression. We have also shown that HBx increases the amount of endogenous OCT3/4 which might improve the process of iPSC generation. It is also possible that HBx affects the cell cycle especially at later stages of the reprogramming process increasing the efficiency iPSC generation. Another possibility is that HBx decreases the apoptosis at various stages of the reprogramming. Significant cell death follows the transfection (“Nucleofection”) of fibroblasts. HBx protein can drive cells division in stressed cells(Kapoor et al., 2013). Reprogramming transcription factors should get access to their specific DNA sequences on the host chromatin. In somatic cells these regions (for example OCT3/4 binding elements) are highly condensed and getting access to the sites is a limiting step in reprogramming. HBx plays an important role in modulating the epigenetic landscape (Kalra and Kumar, 2006; Rivière et al., 2015). It is possible that HBx could improve the efficiency of reprogramming by facilitating the interaction between the reprogramming transcription factors and their binding elements in DNA by loosening up the heterochromatin. HBx is a broad-spectrum activator of transcription; therefore, it might enhance the complex interactions between transcriptional networks decreasing the threshold to establish and sustain the transcriptional networks for the state of pluripotency. HBx is a pleotropic gene, which makes it very difficult to fully elucidate the mechanism of action with reference to reprogramming.

There are concerns regarding the use of *MYC* (a proto-oncogene) in the generation of iPSC. HBx as alluded to earlier is associated with hepatocellular cancer. However, in the iPSC generated using HBx we did not find the expression of HBx protein, nor could we detect HBx integration by PCR. Further, it is possible that HBx may not be as malignant as originally perceived. The HBx protein is known to confer resistance against nucleolar stress(Kapoor et al., 2013),(Kim et al., 2008). Niu Y et al reported that ‘transactivation of FXR by HBx may represent a protective mechanism to inhibit inflammation and the subsequent carcinogenesis’(Niu et al., 2017). It may be noted that HBV viral infection per se is rarely oncogenic in hepatocarcinogenesis(Niu et al., 2017). In either case, since the iPSC generated does not express HBx there is little concern regarding the oncogenic potential of HBx in the iPSC generated.

HBx exerts a clearer effect in high passage fibroblasts could be because of a few reasons. With higher passage, their cell replication capacity decreases. Further, these aged cells are more prone to apoptosis when subjected to the stress of transfection and reprogramming. HBx as shown in Figure 5 C and D improves cells survival upon stress and cell division. This might help reprogramming. It is known that when cells age their chromosomes get more condensed. As discussed before HBx relieves chromatin-mediated transcriptional repression, although we do not know that’s the case here.

Deriving enough cells from small amounts of clinical samples such as fine needle aspiration biopsy (or poor-quality samples) is challenging. Since the starting number of cells are small, they need to undergo more rounds of cell divisions to attain enough numbers for certain experiments. The same is true for cells from higher passages. During continuous culture cells acquire chromosomal abnormalities, deletions, mutations, epigenetic changes and senescence. Here we found high passage (senescent) fibroblast samples producing iPSC colonies when Yamanaka’s factors were supplemented with HBx. The iPSC colonies thus derived were similar (in all aspects we studied) to those which were derived from early passage fibroblasts. This shows that that HBx together with Yamanaka’s factors can probably reverse an ageing (senescent) phenotype although more experiments are required to prove this point.

We have for the first time demonstrated in-vitro that Hbx can increase the cellular levels of OCT3/4. The increased levels of OCT3/4 and MYC is possibly enhancing the generation of iPSC. We have also shown that the iPSC generated using Hbx can be differentiated to all the three germ layers, and terminally to hepatocyte like cells.

Our findings also have some importance in understanding the pathogenesis of hepatocellular carcinoma. HBx might change the epigenetic milieu favoring dedifferentiation and increase in cell division. Both favor the development of cancer as dividing cells are more prone to accumulate mutations over time especially when apoptosis is inhibited.

## Conclusion

Hepatitis B virus x (HBx) protein increases the efficiency of reprogramming of fibroblasts to iPSC. HBx can substitute for MYC protein in the Yamanaka’s factors (OCT4, SOX2, KLF4, MYC). HBx can also improve the generation of iPSC from difficult to reprogram/senescent cell samples using Yamanaka’s factors. This may be helpful for biobanks interested in creating iPSC repositories. Our in-vitro experiments suggest that HBx induces OCT3/4 (and MYC) expression and aids reprogramming. These findings also have implications in understanding the mechanisms of pathogenesis of hepatocellular carcinoma caused by HBV. Further, our findings suggest HBx together with Yamanaka’s factors can probably ‘reverse’ senescence under certain conditions.

## Availability of data and materials/Supplementary data

Madhusudana Girija, Sanal, 2023, “Uncut protein blot (western blot)/DNA agarose gel electrophoresis images”, https://doi.org/10.7910/DVN/NZNDR5, Harvard Dataverse

Any further details or supporting materials can be obtained upon request via email to the corresponding authors.

## Author Contributions

SMG conceived the idea, performed experiments, analyzed, and interpreted results, wrote manuscript, and obtained funds; SG performed experiments-cell culture. RS performed all Western Blots, NV assisted SMG in cloning pHBx, SKS reviewed the manuscript.

## Abbreviations

HBx: Hepatitis B virus X protein
iPSC: induced Pluripotent Stem Cells
YF: Yamanaka’s Factors
HBV: Hepatitis B Virus
cccDNA: covalently closed circular DNA
DMEM: Dulbecco′s Modified Eagle′s Medium
DNA: Deoxyribo Nucleic Acid
PBS: Phosphate Buffered Saline
PEI: Poly Ethylen Imine
TBST: Tris Buffered Saline Tween 20
cDNA: complementary Deoxyribo Nucleic Acid
RT qPCR: Real Time quantitative PCR

## Acknowledgements

We acknowledge Dr. Saurabh Bhattacharya and Dr. Leena Rawai, Lal Path Labs, New Delhi for their help in karyotyping and Dr. Shajo Kunnth, Dr. Anna Velcich and Mr. Alan Alfieri for their critical comments.

## Declarations

- Ethics approval and consent to participate: Authors have taken the required approval from Institutional Ethics Committee/Institutional Review Board for the human samples used.
- Consent for publication: All the authors have agreed upon the content of the manuscript content and the corresponding author on the behalf of all authors consent for publication of the manuscript.
- Availability of data and material: All the required data is available with the corresponding authors and will be submitted to an online public/free repository and made available as per the journal requirement.
- Competing interests: The authors have no competing or conflicting interests to declare.
- Funding: This work was funded partly by two grants to the corresponding author as mentioned under the financial disclosures

---

**Supplementary data Ia.**
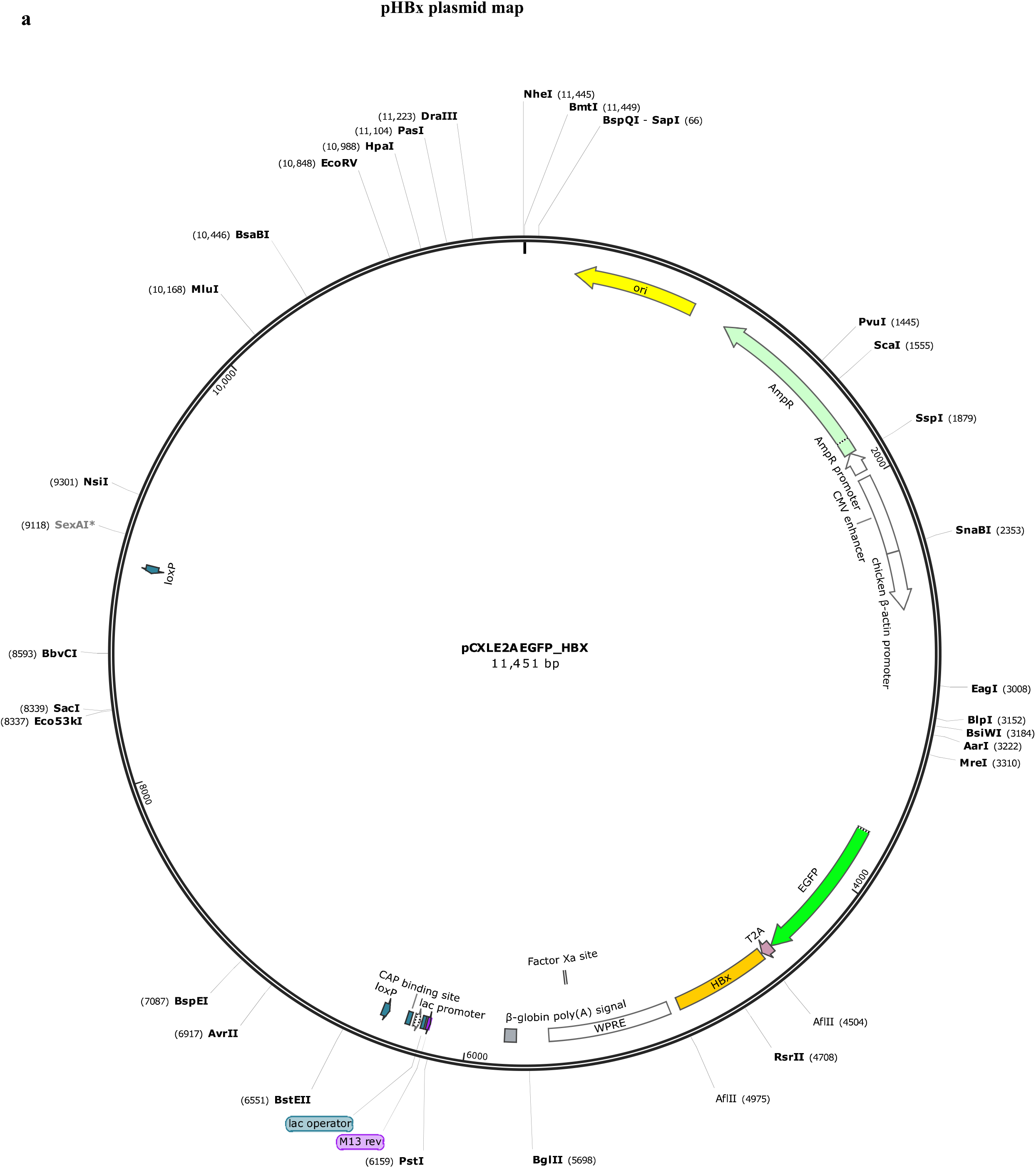
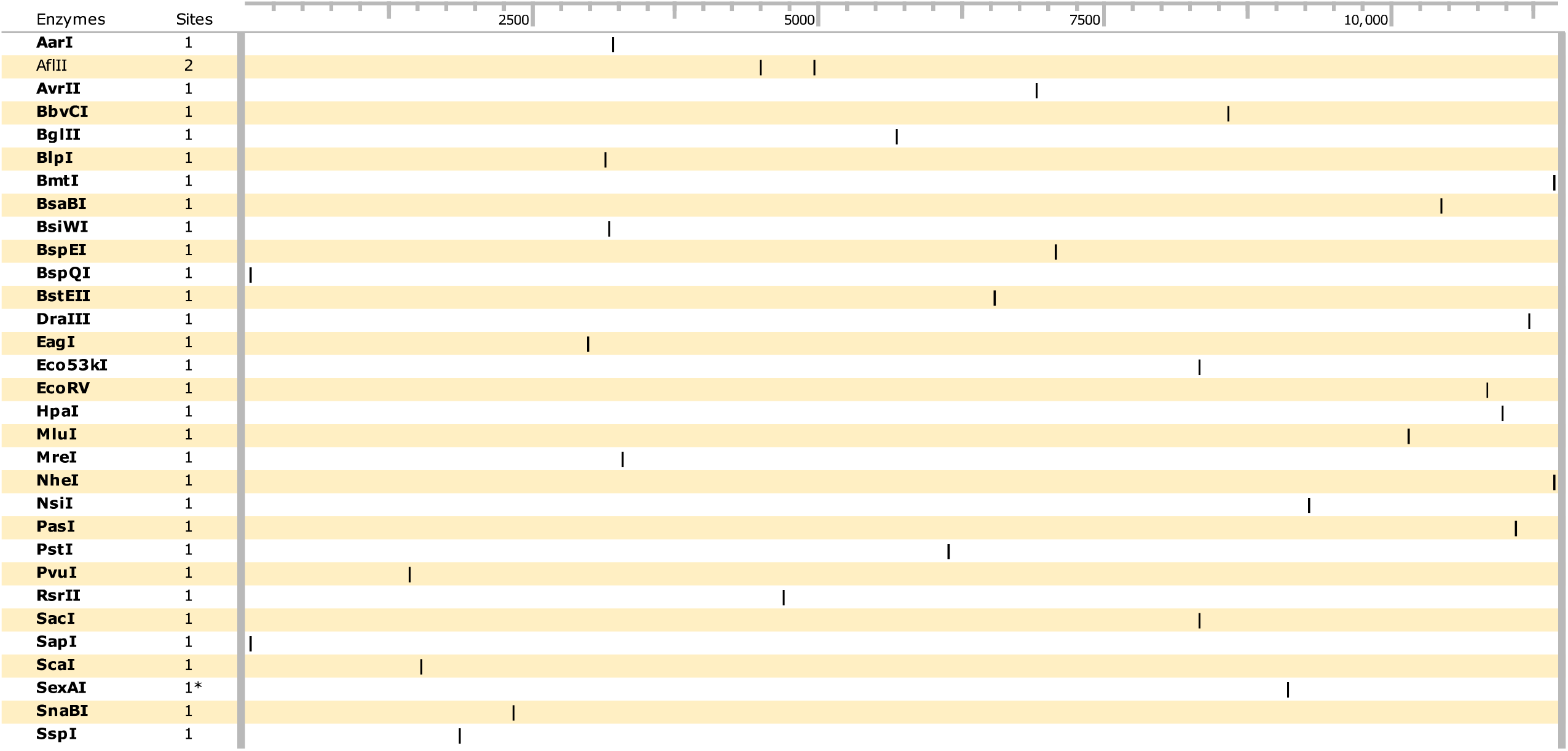

**Supplementary data Ib.**
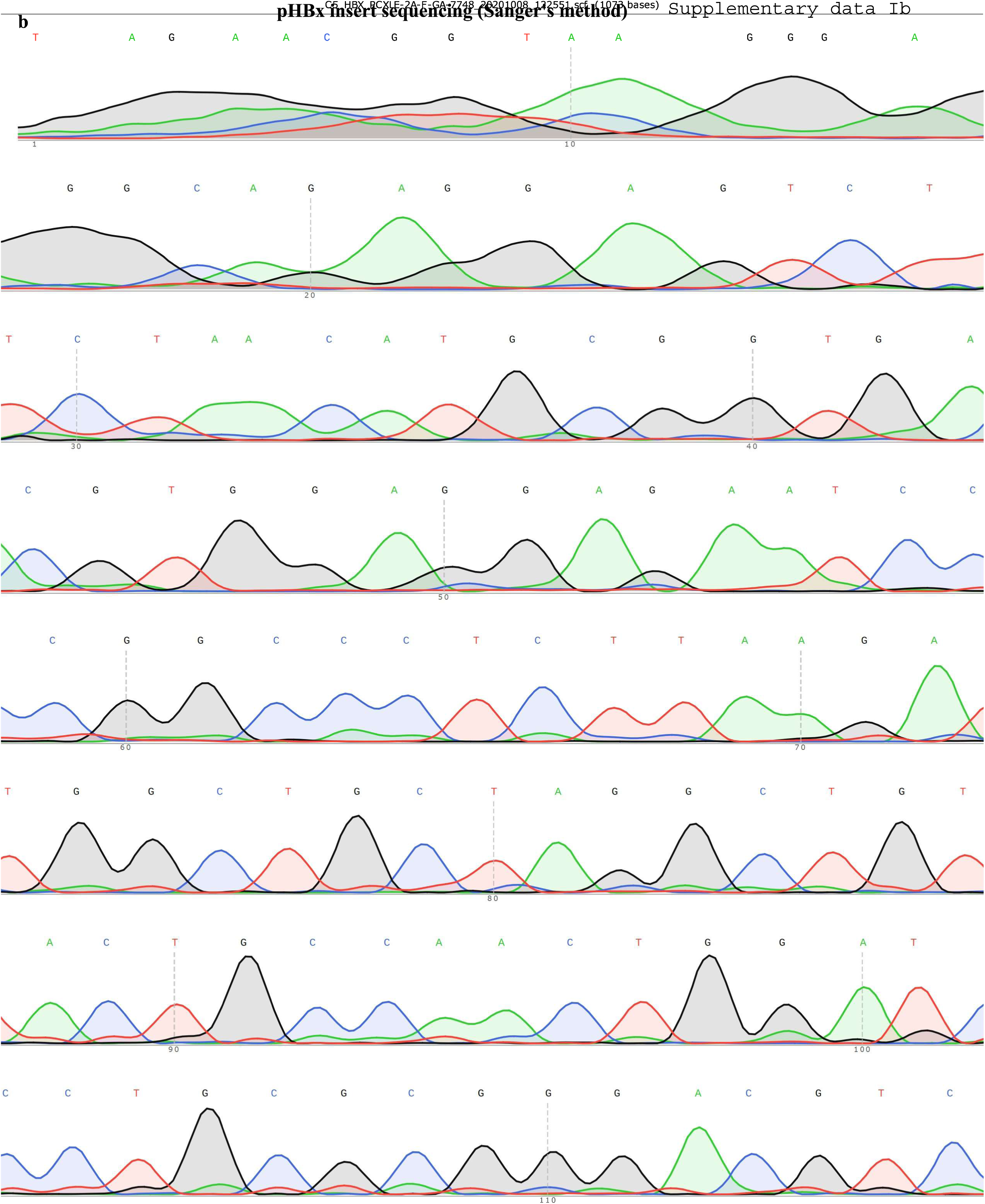
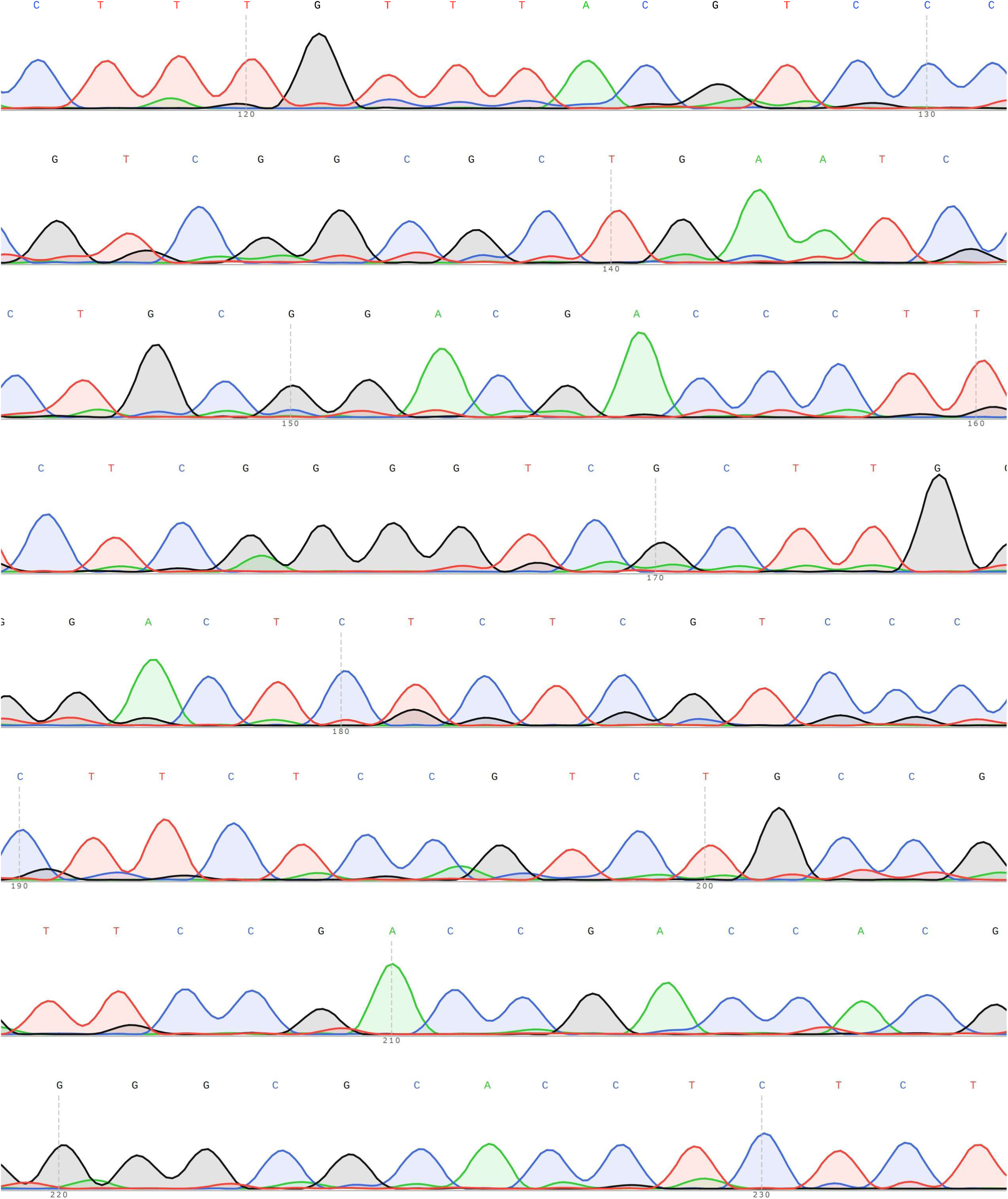
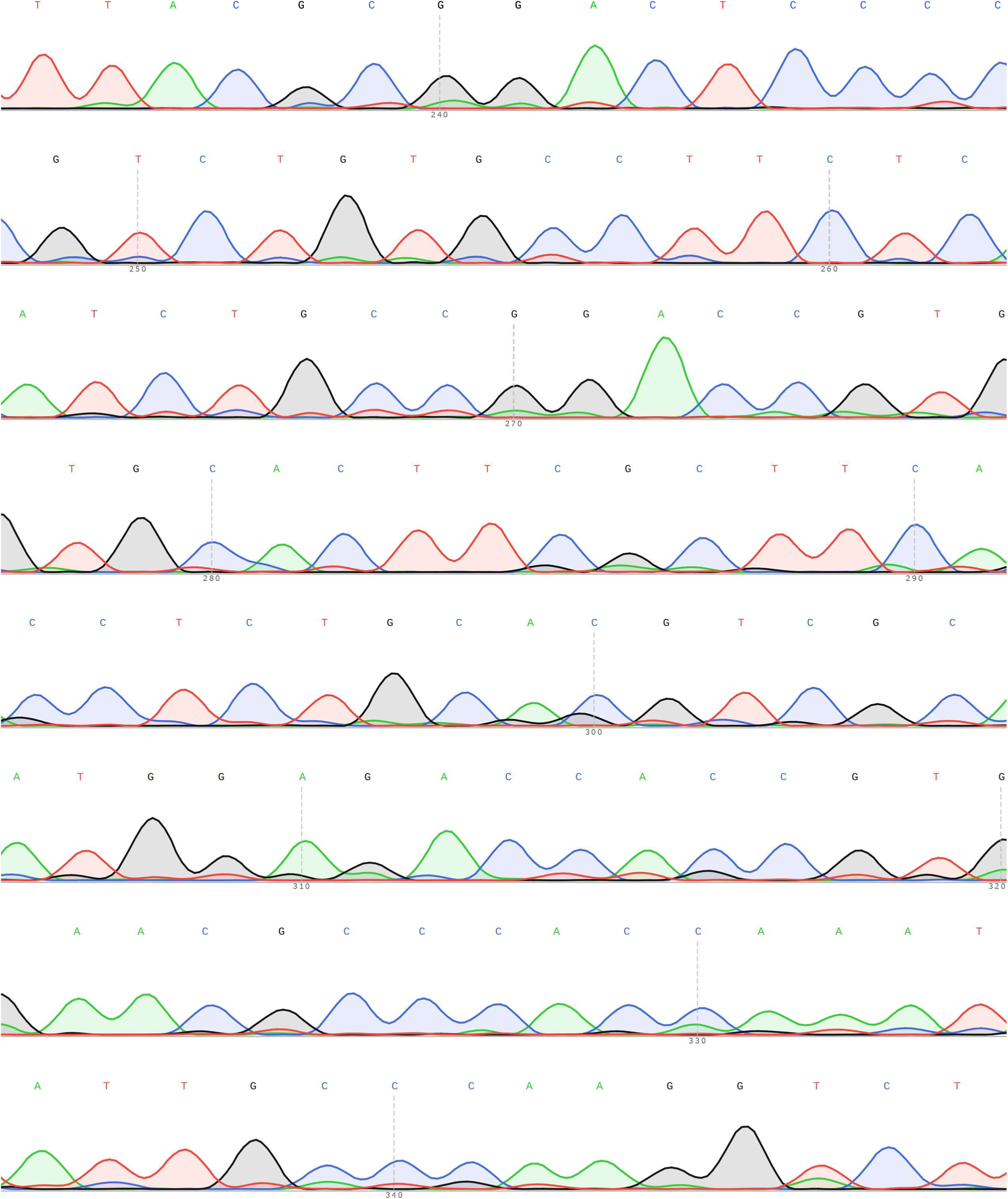
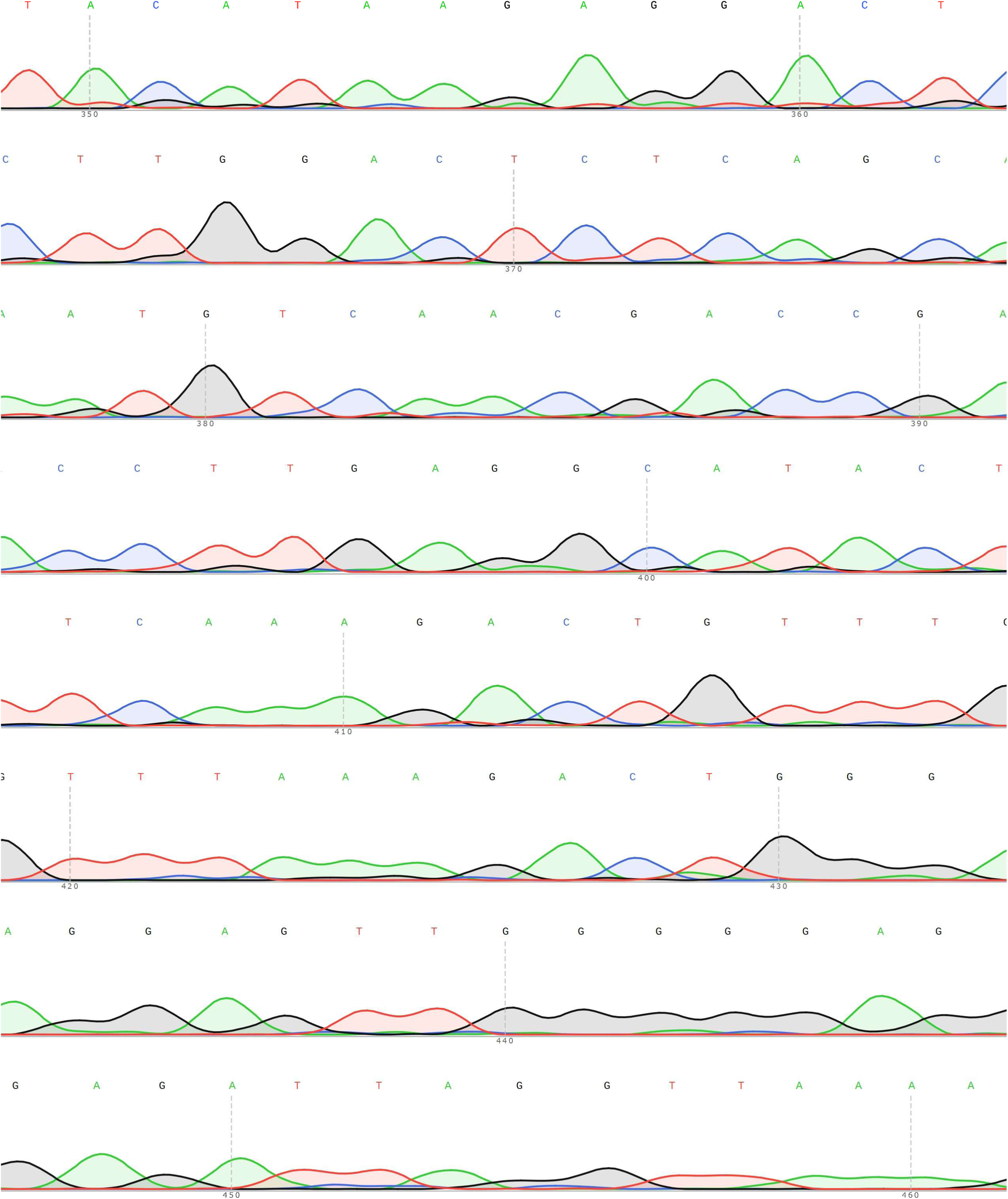

**Supplementary data Ic.**
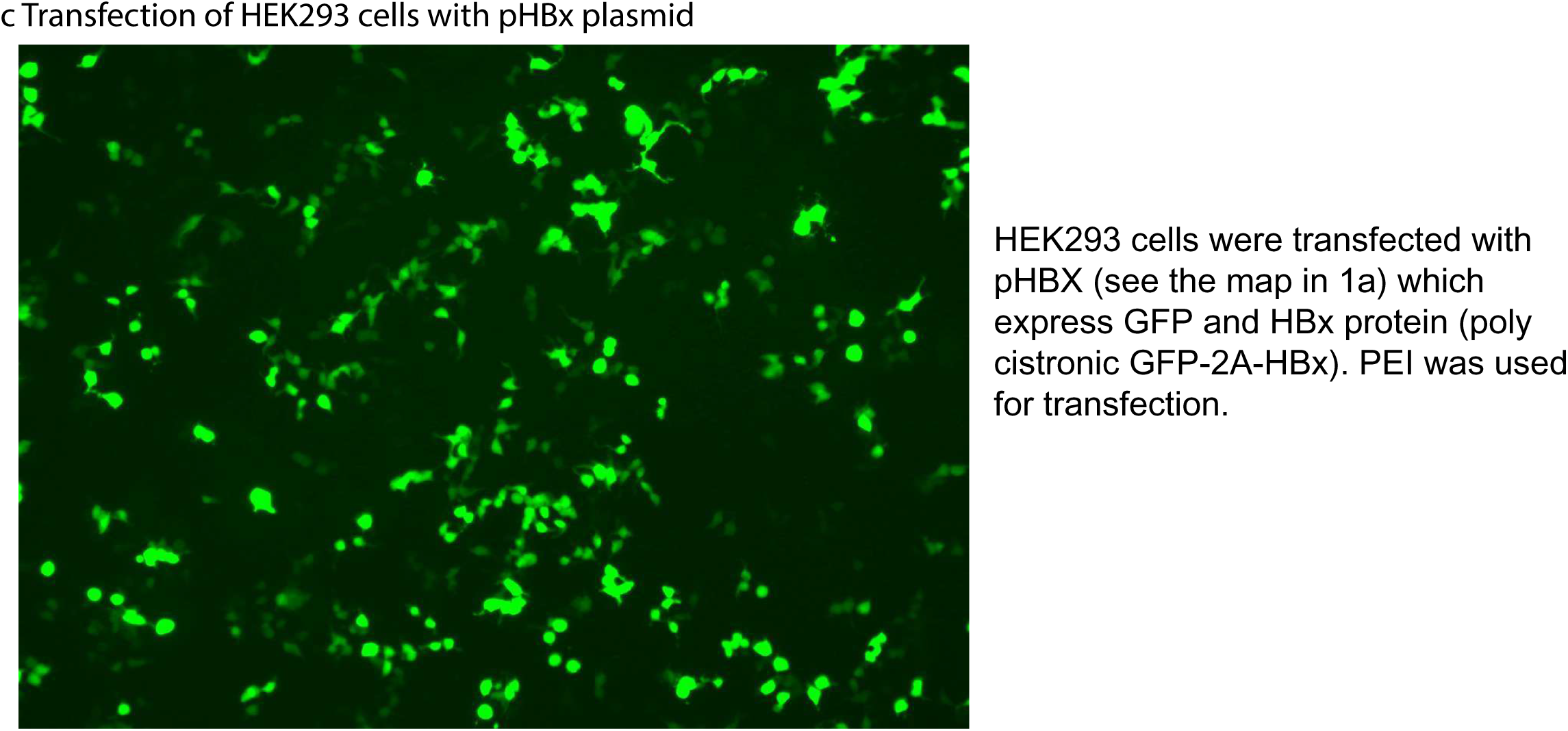

**Supplementary data Id.**
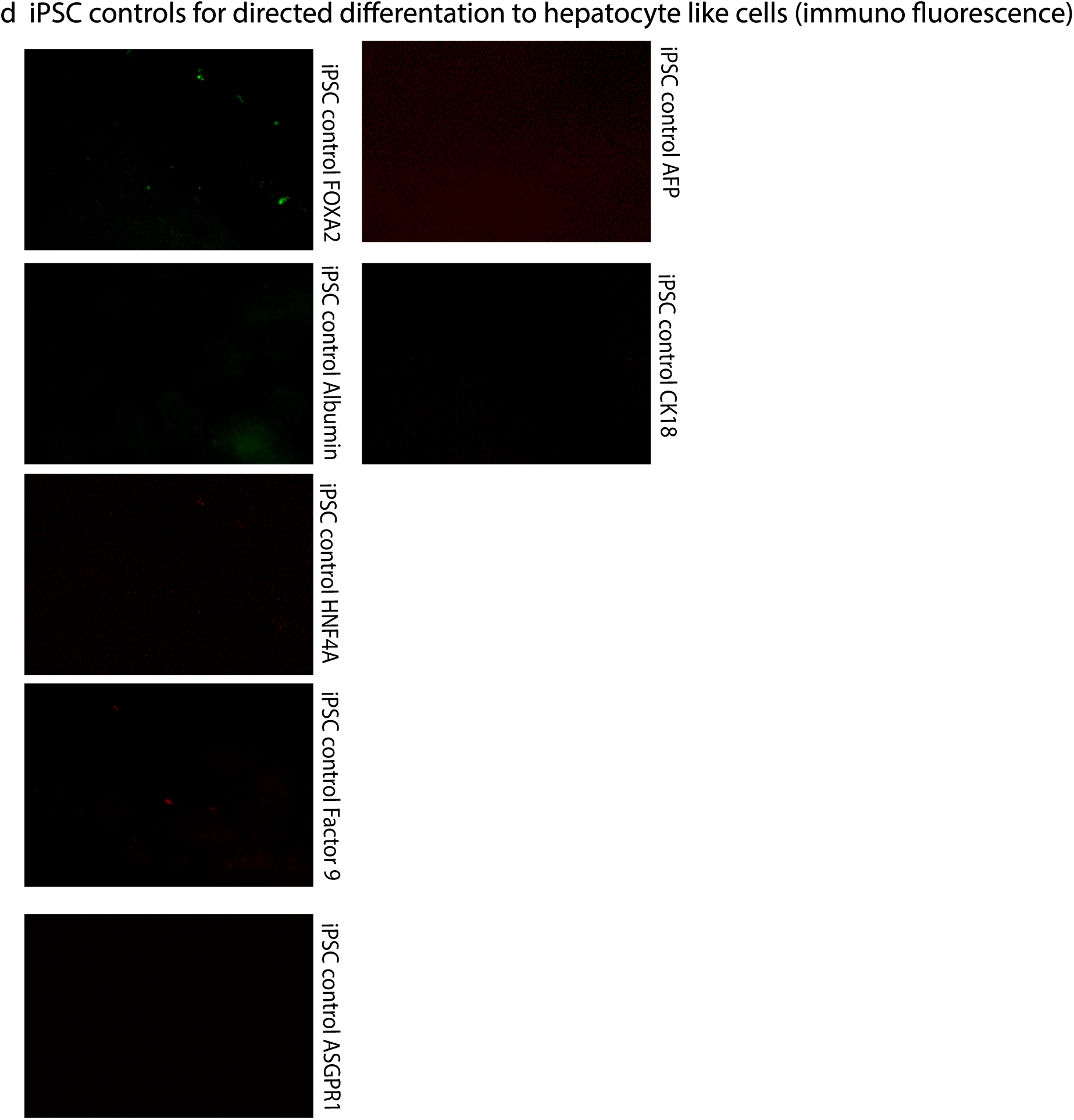

**Table-1:**
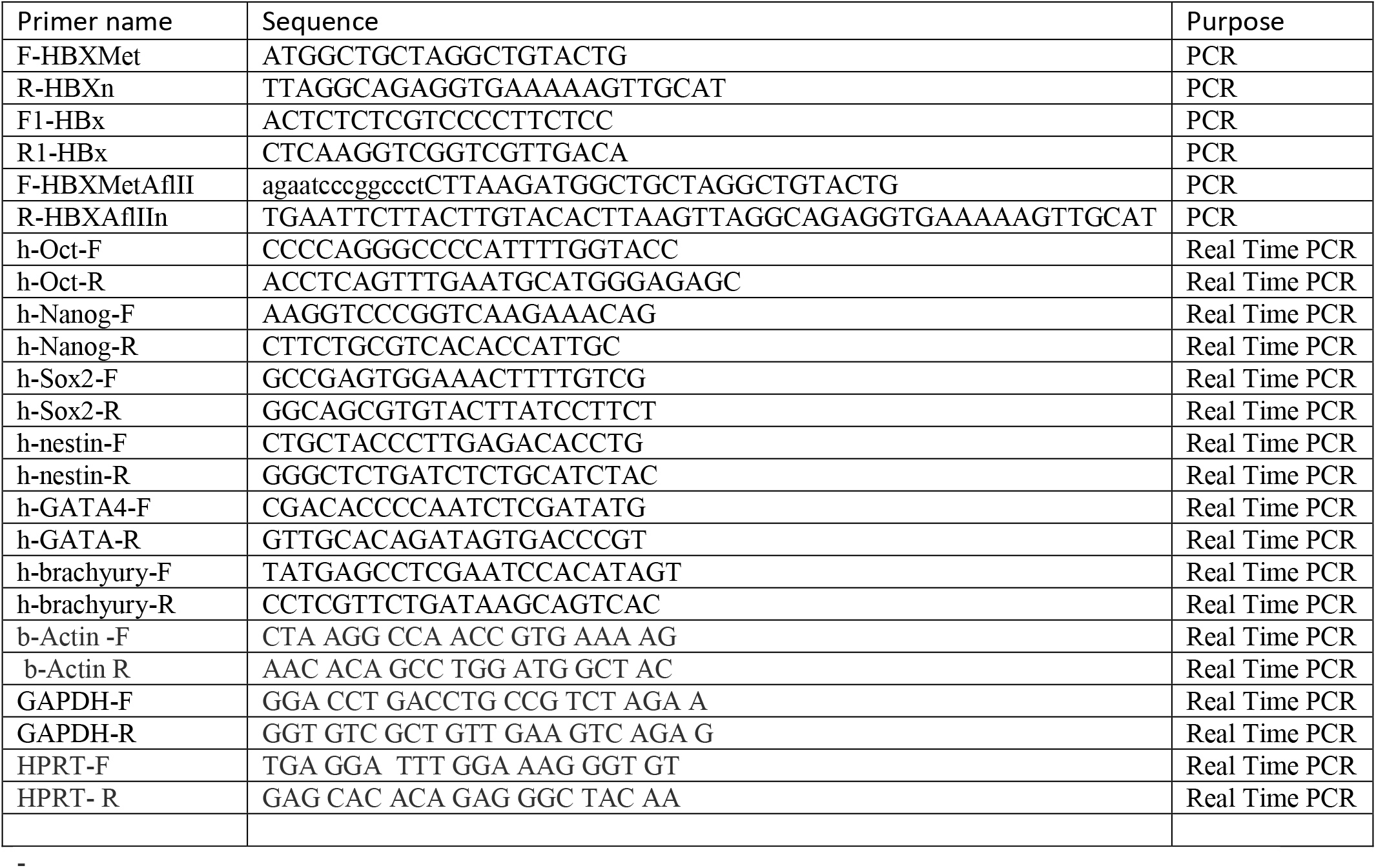
Oligonucleotide primers.

